# Hybridization and diversification are positively correlated across vascular plant families

**DOI:** 10.1101/724377

**Authors:** Nora Mitchell, Kenneth D. Whitney

## Abstract

Hybridization has experimental and observational ties to evolutionary processes and outcomes such as adaptation, speciation, and radiation. Although it has been hypothesized that hybridization and diversification are correlated, this idea remains largely untested empirically. Here, we use a hybridization database on 195 plant families, life history information, and a time-calibrated family-level phylogeny to test for phylogenetically-corrected associations between hybridization and diversification rates, while also taking into account life-history traits that may be correlated with both processes. We show that diversification and hybridization are positively correlated using three different methods to estimate diversification rates and two different metrics of hybridization. Moreover, the relationship remains detectable when incorporating the correlations between diversification and two other life history characteristics, perenniality and woodiness. We further provide potential mechanisms for this association under three different scenarios: hybridization may drive diversification, diversification may drive hybridization, or both hybridization and diversification may jointly be driven by other factors. We suggest future studies to disentangle the causal structure.

## INTRODUCTION

Hybridization is a biological phenomenon whereby different species mate, typically forming offspring that are genetically and often phenotypically distinctive. The prevalence of natural hybridization is becoming increasingly recognized in animals (involving up to 25% of species in some groups (Mallet 2005), fungi (reviewed in Albertin and Marullo 2012), and in plants, with hybridization occurring in 40% of plant families (Whitney et al. 2010a) and involving up to 25% of species (Mallet 2005).

Hybridization increases the availability of genetic variation on which natural selection can act (Schluter et al. 2004; Barrett and Schluter 2008) and can result in novel phenotypic traits or in novel combinations of traits, e.g. via transgressive segregation (Rieseberg et al. 1999; Bell and Travis 2005; Dittrich-Reed and Fitzpatrick 2013). Moreover, introgressive hybridization, or repeated backcrossing from one lineage into another, can result in the transfer of adaptive traits between lineages that confer advantages in novel environments, explored in (Anderson 1953; Dowling and Secor 1997; Suarez-Gonzalez et al. 2018).

Both theory and empirical observations support the idea that hybridization can promote increased rates of evolution and/or speciation (Anderson and Stebbins 1954; Stelkens et al. 2014; Marques et al. 2019; Taylor and Larson 2019). Hybridization has been linked to multiple evolutionary and ecological processes, such as adaptation, e.g. (Lewontin and Birch 1966; Campbell and Snow 2007; Hovick et al. 2012; Stankowski and Streisfeld 2015; Mitchell et al. 2019a), speciation, e.g. (Rieseberg 2003; Mallet 2007; Rieseberg et al. 2007; Soltis and Soltis 2009; Abbott et al. 2013), and evolutionary radiation, e.g. (Anderson and Stebbins 1954; Stebbins 1959; Barton 2001; Seehausen 2004; Yakimowski and Rieseberg 2014; Berner and Salzburger 2015; Marques et al. 2019). For example, on a microevolutionary scale, hybridization has resulted in the adaptive introgression of traits related to both abiotic and biotic factors in Texas sunflowers (*Helianthus annuus* ssp. *texanus*) (Whitney et al. 2006, 2010b). Experimental hybridization in this system increased the speed of adaptive evolution over eight generations when compared to non-hybrid controls (Mitchell et al. 2019a). Hybridization has also led to speciation within this same genus. Historical interbreeding between two sunflowers (*Helianthus annuus* and *H. petiolaris*) generated three species of hybrid origin that are adapted to novel and extreme environments in a repeatable fashion (reviewed in Rieseberg et al. 2007). On a macroevolutionary scale, hybridization has been linked to rapid speciation and evolutionary radiations. For example, in the Hawaiian silverswords, an ancient hybrid founder may have provided the evolutionary novelty necessary to promote adaptive radiation (Barrier et al. 1999).

Based on the association between hybridization and evolutionary change at different scales, it has been hypothesized that hybridization may be linked to overall net diversification (Dowling and Secor 1997; Seehausen 2004), yet this idea remains untested (but see Tank et al. 2015; Landis et al. 2018 for related work linking polyploidy to diversification). Importantly, the direction of causality between hybridization and diversification could go either way; high rates of hybridization could result in high rates of diversification, or high rates of diversification could result in the increased prevalence of hybridization within a lineage. For instance, in the latter scenario, high rates of diversification may mean that species have low amounts of genetic divergence between them, allowing for higher rates of hybridization.

Net diversification, the collective result of speciation minus extinction, can be estimated from time-calibrated phylogenies (Ricklefs 2007). Diversification is an emergent property of a given lineage and is influenced by individual, population, or species-specific factors that can increase or decrease the likelihood of either speciation or extinction (Barraclough 1998; Langerhans and DeWitt 2004; Bouchenak-Khelladi et al. 2015). It is therefore necessary to investigate the relative effects of different factors on diversification in a multivariate context, while acknowledging that additional yet unmeasured factors (and their interactive effects) are likely at play and that the explanatory power of any one factor is likely small.

Here, we present a test of the hypothesized correlation between hybridization and diversification in vascular plants. Both diversity (Davies et al. 2004) and rates of hybridization (Ellstrand et al. 1996; Whitney et al. 2010a; Beddows and Rose 2018; Mitchell et al. in 2019b) are unevenly distributed across plant lineages. Moreover, both diversification and hybridization have been linked to numerous aspects of plant biology, including life form and life history. For instance, plant groups dominated by herbaceous life forms or short generation times tend to have faster diversification rates (Eriksson and Bremer 1992; Dodd et al. 1999), while both perenniality and woodiness are positively associated with hybridization (Ellstrand et al. 1996; Beddows and Rose 2018; Mitchell et al. 2019b).

Using a database containing two measures of hybridization and life history information on 195 vascular plant families, as well as a time-calibrated family-level phylogeny, we estimate family-level diversification rates using three different methods and ask 1) Are diversification rates and hybridization rates related across vascular plants?, and 2) Does hybridization remain predictive of diversification when accounting for other correlated traits?

## MATERIALS AND METHODS

### Hybridization rates, perenniality, and woodiness

For hybridization rates in vascular plant families, as well as two life history characteristics (perenniality and woodiness) known to be positively correlated with hybridization rates (Mitchell et al. 2019b), we used the hybrid and trait database previously examined in Mitchell et al. (2019b). This database includes data from eight regional floras: the Great Plains of the U.S. (McGregor and Barkley 1986) the British Isles (Stace 1997); Hawai’i (Wagner et al. 1999); the Intermountain Region of the western U.S. (Cronquist et al. 1972); the Northeastern U.S. (Magee and Ahles 1999); California (Hickman 1993); Europe (Tutin et al. 1964); and Victoria, Australia (Walsh and Entwisle 1994). In summary, for each plant family in each flora the number of non-hybrids and interspecific hybrids was assessed as in Whitney et al. (2010). For counting purposes, a “hybrid” was defined as a hybrid type derived from a unique combination of two parental species (as in Ellstrand et al. 1996). Thus, in each flora, each pair of hybridizing species was counted as generating a single hybrid, even if there was evidence that the pair had hybridized multiple times. Recognition of an interspecific hybrid does not imply that it was formally or taxonomically recognized in the flora (though some were). From these data, two metrics of hybridization (hybridization propensity and hybrid ratio were estimated). Hybridization propensity (HybProp) is calculated as

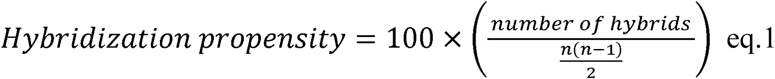

where *n* = the number of nonhybrid species in the family. It thus represents the percentage of possible hybrid combinations that have been actually realized in nature. Hybrid ratio (HybRatio) is calculated as

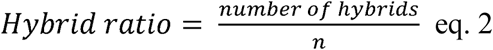

and has been used in previous studies examining patterns of hybridization (e.g. Mitchell et al. 2019b; Beddows and Rose 2018). Note the scale difference: by convention, hybridization propensity is a percentage bounded between 0 and 100, while hybrid ratio is unbounded (in the Mitchell et al. 2019b dataset, it ranges from 0 – 0.15 with outliers up to 1.2). Overall, the dataset contains hybridization data on 282 plant families, and after eliminating genera where a single non-hybrid species was observed in a single flora (thus having no opportunity for hybridization), 195 plant families were retained for subsequent analyses. See Mitchell et al. (2019b) for details on collection of these data.

We used the values of perenniality and woodiness estimated by Mitchell et al. (2019b). Briefly, the number of annual, biennial, and perennial species, and the number of herbaceous vs. woody species were counted in the floras. Each species was assigned a score (for perenniality: 0 if annual, 0.5 if biennial, 1 if perennial; for woodiness: 0 if herbaceous, 1 if characterized by aboveground woody biomass; with intermediates split between categories). Perenniality and woodiness were then estimated as percentages of species within a family possessing the characteristic.

### Vascular plant phylogeny

We used the family-level seed plant phylogeny of Qian and Zhang (2014) to assess diversification rates (while incorporating family-level species richness, see below) and account for phylogenetic non-independence of lineages. Briefly, the Qian and Zhang (2014) phylogeny was built by extracting a single species from each family from the Zanne et al. (2014) species-level phylogeny, updating the taxonomy, correcting errors, and including six additional families to produce a phylogeny with tips representing 437 seed plant families with branch lengths proportional to time in millions of years. The phylogeny was downloaded from Appendix S3 of Qian and Zhang (2014). We checked taxonomy according to The Plant List (<www.theplantlist.org>, as of March 2019) and trimmed families from the phylogeny that are no longer recognized, resulting in a seed plant phylogeny of 413 families. To incorporate ferns and their allies, we used the time-calibrated phylogeny of Testo and Sundue (2016). We trimmed the fern phylogeny to a random species from each family and bound the seed plant phylogeny to the family-level fern phylogeny. We then ultrametricized the product using the force.ultrametric() function and the “nnls” method in the R package phytools (Revell 2012), resulting in a vascular plant phylogeny with 459 families. The phylogeny and additional metrics were visualized using the R package ggtree (Yu et al. 2017).

### Species richness

We created a taxonomic richness matrix based on the number of species in each family listed in The Plant List (<www.theplantlist.org>, as of March 2019).

### Diversification rates

We used the family-level phylogeny to assess diversification rates using three different methods: Method-of-moments estimation, Medusa, and BAMM.

#### Method-of-moments (MS)

We used the stem-group method-of-moments estimation (MS) of diversification using the stem ages of each family (tip edge lengths from the family-level phylogeny) and family species richness to calculate a measure of net diversification rates (Rohatgi 1976; Magallon and Sanderson 2001). We calculated MS using three different relative extinction rates (epsilon, □ = 0, 0.5, or 0.9) and the equation *r* = ln(*n* (1 - □) + □)/*t*, where *r* = the net diversification rate, *n* = the extant species richness and *t* = the stem group age (in millions of years).

#### Medusa

We used Medusa (Alfaro et al. 2009) to model diversification using stepwise AIC using piecewise birth-death models. We used the medusa() function in the R package geiger (Harmon et al. 2008) using the family-level phylogeny and family species richness. The program compares stepwise models until the improvement, measured using the sample-size corrected Akaike information criterion (AICc), does not exceed the internally computed threshold (for a phylogeny with 459 tips, threshold = 8.089111). Because results are highly dependent on threshold value (May and Moore 2016), we ran Medusa using several threshold values (8.089, 4, and 2) to compare outputs; note that a previous version of Medusa used a default threshold value of 4. We extracted the net diversification rates associated with each tip (family) from each of these threshold runs.

#### BAMM

We used BAMM (Bayesian Analysis of Macroevolutionary Mixture Models) (Rabosky 2014) to model speciation and extinction rates using reversible jump Markov chain Monte Carlo to explore model space. Prior values were obtained using the BAMMtools R package (Rabosky et al. 2014). We tested four values for the expected number of shifts (25, 50, 100, and 200), and accounted for richness by providing a sampling fraction for each family calculated as 1 over the number of species in the family. To account for the fact that the “sampling fraction” was often extremely low, we set extinctionProbMax = 0.9999999. We ran BAMM on the family-level seed plant phylogeny on the high-performance computing facilities maintained by the UNM Center for Advanced Research Computing for 50 million generations sampled every thousand generations. We processed the output in BAMMtools and used a burnin of 20%, leaving 40,001 samples. Convergence was checked visually using the R package coda (Plummer et al. 2006); we ensured that the effective sample sizes for the number of shifts and log-likelihoods were over 200. We then extracted net diversification rates for each family.

### Phylogenetic Generalized Least Squares (PGLS)

We used phylogenetic generalized least squares regression (PGLS) (Grafen 1989; Martins and Garland Jr 1991) to detect associations between diversification rates and rates of hybridization in plant families while accounting for evolutionary history. We trimmed the family-level phylogeny to include only the families for which we had hybridization data, resulting in a phylogeny of 195 families. We standardized all variables to make results comparable between analyses. We ran univariate PGLS using the pgls() function in the R package caper (Orme et al. 2013), modeling diversification rates as outcomes and using each hybridization metric as our predictor with estimated lambda values and kappa and delta set to 1.

For each combination of diversification and hybridization metrics, we also ran multivariate regressions using standardized data to detect associations between diversification rates and hybridization metrics when including two other predictors: perenniality and woodiness.

### Sister-clade comparison

We further examined the relationship between hybridization and diversification using the more traditional approach of sister-clade comparisons (Slowinski and Guyer 1993; Barraclough 1998) to ask whether families with higher rates of hybridization also had higher diversification rates. We identified sister families present in the 459-taxon phylogeny and consistent with relationships on the Angiosperm Phylogeny Website <http://www.mobot.org/MOBOT/research/APweb/, accessed 23 April 2019>. We restricted comparisons to sister pairs for which we had data and where both hybridization propensity and hybridization ratio differed consistently (i.e., the same family had higher values for both metrics), as there were some cases where the metrics were inconsistent in identifying the more hybridization-prone family. This resulted in 11 sister-clade comparisons. We then asked whether the diversification rates differed between families in the same direction that hybridization differed using a non-parametric one-tailed sign-test.

## RESULTS

### Hybridization rates

Hybridization rates across families in this dataset have been presented and discussed previously in Mitchell et al. (2019b). Briefly, out of 195 plant families, 112 contained hybrids. The average family-level hybridization propensity was 2.55 and the average hybridization ratio was 0.086. See Mitchell et al. (2019b) Fig. 2 and Table S2 for details.

### Diversification rates

We used three different methods to estimate diversification rates across the family-level vascular plant phylogeny: method-of-moments (MS), Medusa, and BAMM.

We estimated the MS net diversification rates using relative extinction fractions of □ = 0.0, 0.5, and 0.9. Although we are unable to use model comparison since the response measures are different, diversification rates were highly correlated across values of □ (Table S1). We chose to examine rates computed with the moderate value of □ = 0.5 for presentation in the main analyses. The mean net diversification rate was 0.050 species per Ma (range = 0.000 – 0.255) across all 459 families in the phylogeny; the mean rate was 0.077 species per Ma across the 195 families for which we had hybridization data.

Using the computed AICc threshold value of 8.089 for our 459-taxon phylogeny, Medusa detected 48 rate shifts in diversification, while the previously standard threshold value of four detected 82 rate shifts and a threshold of two detected 104 rate shifts. Rate shifts for Medusa runs using different thresholds were highly correlated (Table S2). We chose to examine rates computed using the threshold of four for presentation in the main results. The mean net diversification rate was 0.072 species per Ma (range = 0.000 – 0.280) across all 459 families in the phylogeny and the mean rate was 0.088 species per Ma across the 195 families for which we had hybridization data.

The number of rate shifts detected by BAMM was largely robust to the prior on the expected number of shifts, where the priors of 25, 50, 100, and 200 resulted in 74.2, 77.5, 79.8, and 79.8 mean rate shifts, respectively (Fig. S1). The diversification rates estimated using the four different priors are highly correlated with each other (Table S3). We chose to examine rates estimated using a prior of 100 expected shifts for presentation in the main results, as the posterior estimate was consistent between values of 100 and 200 and differed slightly between 50 and 100. The mean net diversification rate was 0.044 species per Ma (range = −0.069 – 0.162) across all 459 families in the phylogeny and the mean rate was 0.057 species per Ma across the 195 families for which we had hybridization data.

Net diversification rates across the three different methods were highly correlated across the full 459-taxon dataset (MS – Medusa: r = 0.808, p = 0.000; MS – BAMM: r = 0.817, p = 0.000; Medusa – BAMM: r = 0.878, p = 0.000) (Figs. 1, 2).

**Figure 1.**
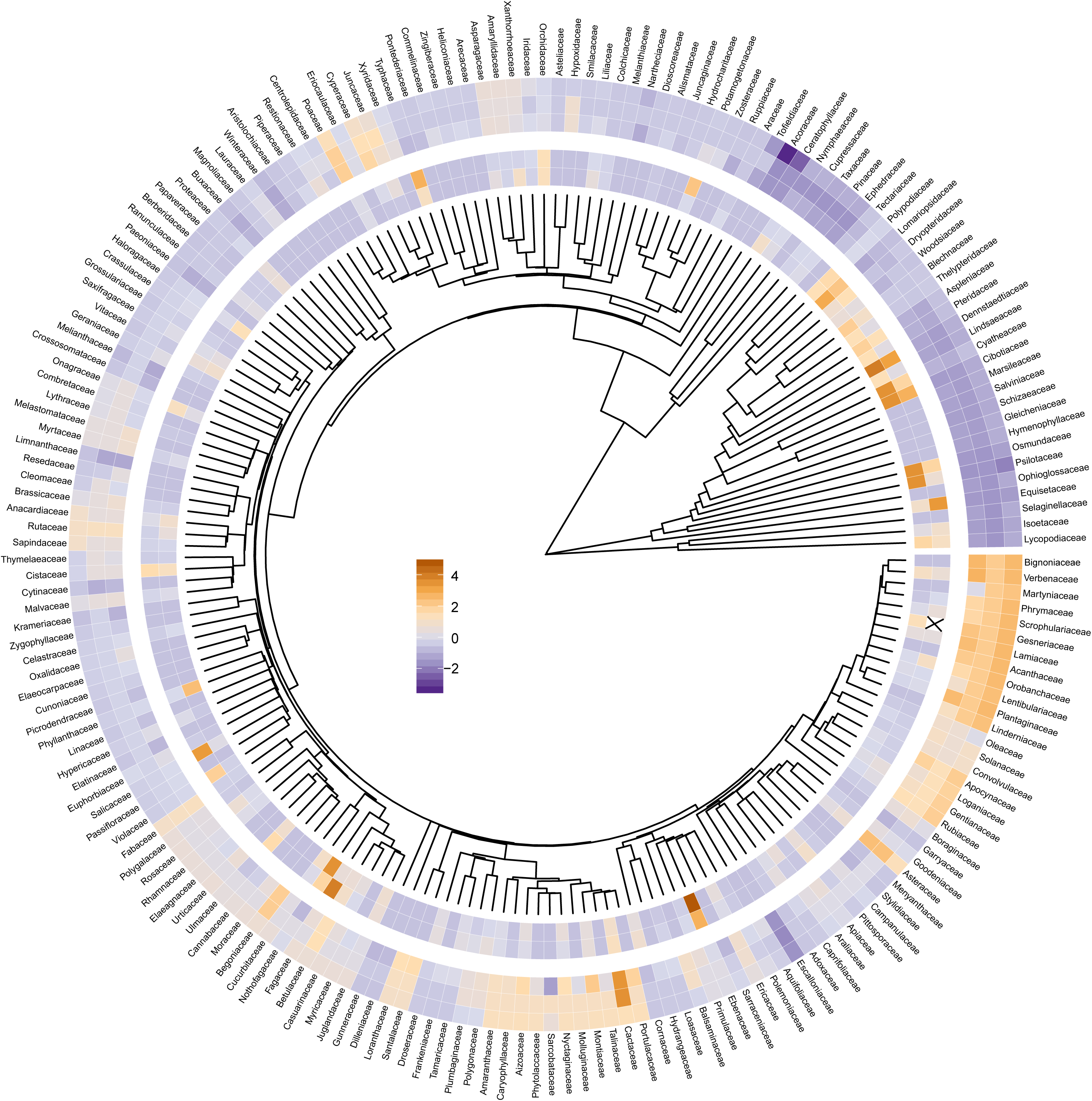
Phylogeny containing 195 vascular plant families analyzed in this study. Rings depict the standardized values of the log-transformed rates for (moving from inside to outside) the hybridization measures (HybProp and HybRatio) and the net diversification rates (MS, Medusa, and BAMM). Phylogenetically corrected analyses typically find a positive association between hybridization and diversification (Fig. 3, Tables 1 and 2), but non-phylogenetically corrected analyses find no relationship (Table S8). Note that the value for the standardized log-transformed HybRatio in Genseriaceae was very high (5.953) and was not plotted (replaced with an “X”) to improve readability of the remaining data. Blue = relatively slow diversification rates / low hybridization rates, orange = relatively fast diversification rates / high hybridization rates.

**Figure 2.**
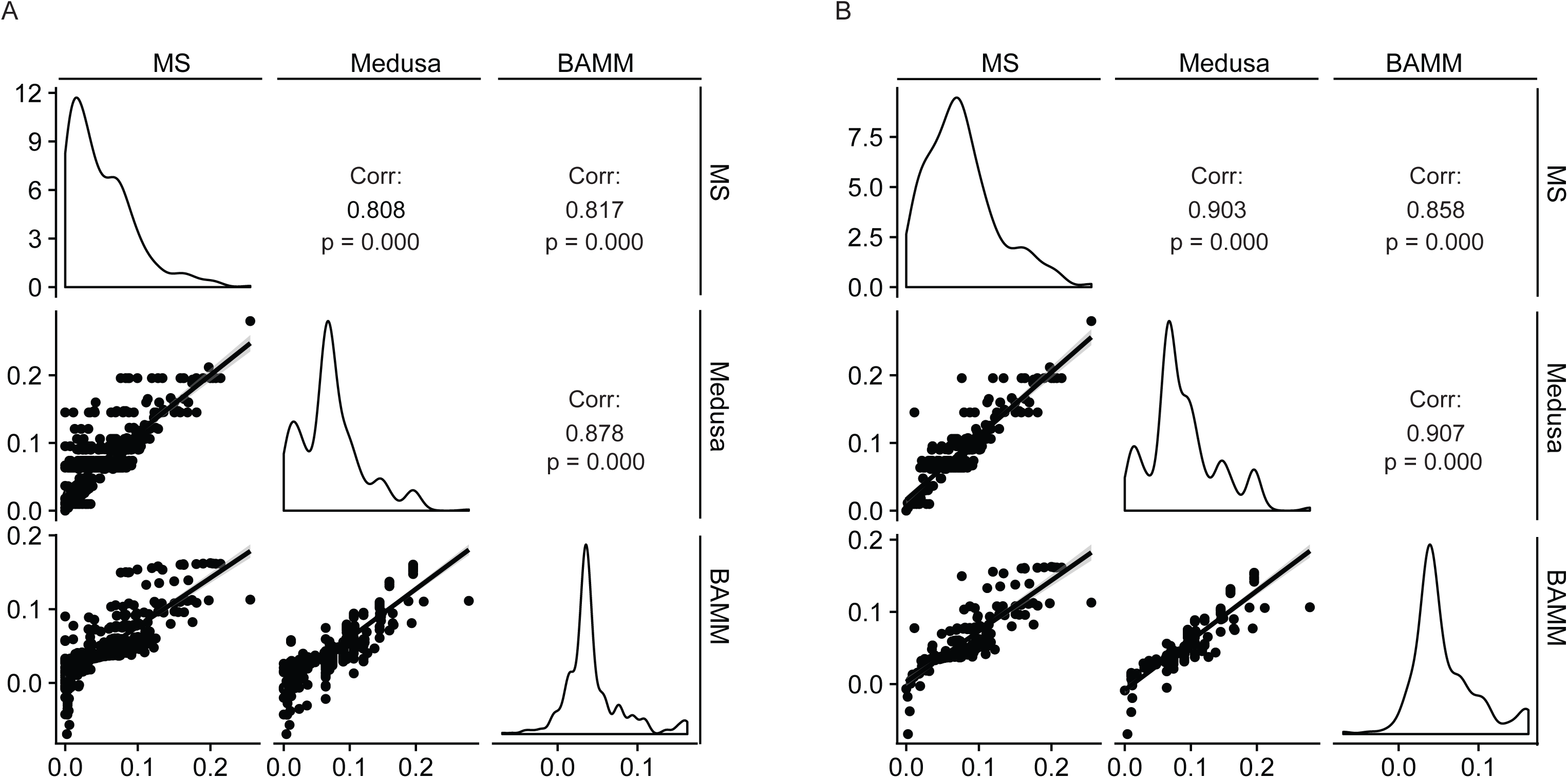
Raw net diversification rates calculated using three different methods are correlated for (A) the full 459 family dataset and (B) the subset 195 family dataset.

### Some evidence for phylogenetic correlation between hybridization and diversification

Univariate phylogenetically generalized least squares (PGLS) regressions of hybridization measures on diversification rates were all positive, meaning that families with higher hybridization rates also had higher diversification rates when accounting for phylogenetic nonindependence (Table 1, Fig. 3A). Using HybProp, hybridization was weakly associated (0.05 < p < 0.10) with increased diversification rates estimated using all three methods. Using HybRatio, hybridization was positively associated with the MS and Medusa diversification rates (p < 0.05) and hybridization was weakly associated with the BAMM diversification rates (0.05 < p < 0.10) (Table 1, Fig. 3A). The relationships were weak (see adjusted-R^2^ values in Table 1), though the results were largely robust to the use of alternate parameters, thresholds, or priors for estimating diversification rates (see Tables S4, S5, S6 for details of supplementary modeling results).

**Table 1.** Univariate PGLS modeling results.

| Method | Hybrid Metric | Adj-R<sup>2</sup> | Estimate | Std. Error | T-value | P-value |
| --- | --- | --- | --- | --- | --- | --- |
| MS | HybProp | 0.011 | 0.116 | 0.066 | 1.754 | 0.081 |
|  | HybRatio | 0.023 | 0.135 | 0.057 | 2.372 | 0.019 |
| Medusa | HybProp | 0.013 | 0.102 | 0.055 | 1.859 | 0.065 |
|  | HybRatio | 0.016 | 0.090 | 0.044 | 2.021 | 0.045 |
| BAMM | HybProp | 0.009 | 0.080 | 0.047 | 1.691 | 0.093 |
|  | HybRatio | 0.012 | 0.066 | 0.036 | 1.817 | 0.071 |

**Figure 3.**
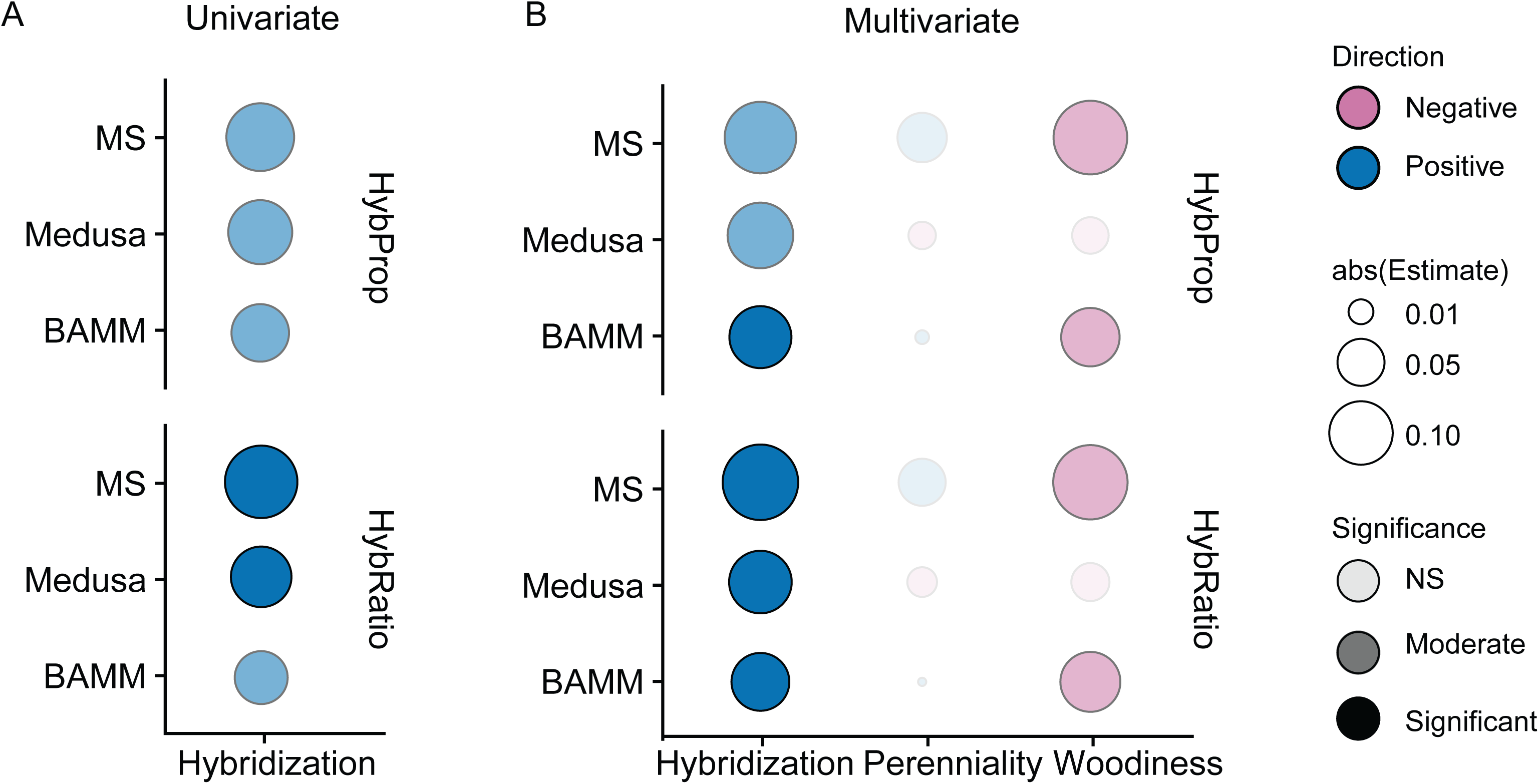
PGLS results for (A) univariate PGLS regressions or (B) multivariate PGLS regressions between either hybridization propensity or hybrid ratio and net diversification as estimated by MS, medusa, or BAMM. Multivariate regressions include the percent of species within each family that are perennial (Perenniality) and that are woody (Woodiness) as additional covariates. Circle size corresponds to the absolute value of the partial regression coefficient, color indicates the direction of the relationship (blue = positive, pink = negative) and transparency indicates the signficance of the relationship (lightest = not significant, medium = 0.05 < p < 0.10, and dark = p < 0.05).

### Correlations between hybridization and diversification are not driven by perenniality or woodiness

Multivariate PGLS regressions incorporating family-level perenniality and woodiness measures reinforced the trends observed in the univariate analyses. Relationships between diversification rates and hybridization measures were consistently positive, though the models overall had very low R^2^ values (Table 2). In the combination of HybProp and BAMM-estimated diversification, the hybridization – diversification relationship was positive and significant at p < 0.05; but when diversification was estimated by MS or Medusa, the hybridization – diversification relationships were weaker but still trended positive (0.05 < p < 0.10). All three diversification measures were significantly positively associated with HybRatio (p < 0.05) (Table 2, Fig. 3B).

**Table 2.** Multivariate PGLS modeling results.

| Method | Hybrid Metric | Adj-R<sup>2</sup> | Covariate | Estimate | Std. Error | T-value | P-value |
| --- | --- | --- | --- | --- | --- | --- | --- |
| MS | HybProp | 0.019 | Hybridization | 0.131 | 0.067 | 1.949 | 0.053 |
|  |  |  | Perenniality | 0.056 | 0.067 | 0.844 | 0.400 |
|  |  |  | Woodiness | -0.140 | 0.074 | -1.891 | 0.060 |
|  | HybRatio | 0.033 | Hybridization | 0.148 | 0.058 | 2.573 | 0.011 |
|  |  |  | Perenniality | 0.050 | 0.066 | 0.748 | 0.455 |
|  |  |  | Woodiness | -0.143 | 0.073 | -1.949 | 0.053 |
| Medusa | HybProp | 0.005 | Hybridization | 0.108 | 0.056 | 1.932 | 0.055 |
|  |  |  | Perenniality | -0.013 | 0.053 | -0.254 | 0.800 |
|  |  |  | Woodiness | -0.027 | 0.060 | -0.450 | 0.653 |
|  | HybRatio | 0.009 | Hybridization | 0.097 | 0.046 | 2.123 | 0.035 |
|  |  |  | Perenniality | -0.016 | 0.053 | -0.303 | 0.762 |
|  |  |  | Woodiness | -0.030 | 0.060 | -0.499 | 0.619 |
| BAMM | HybProp | 0.018 | Hybridization | 0.095 | 0.048 | 1.981 | 0.049 |
|  |  |  | Perenniality | 0.002 | 0.043 | 0.055 | 0.956 |
|  |  |  | Woodiness | -0.083 | 0.050 | -1.665 | 0.098 |
|  | HybRatio | 0.022 | Hybridization | 0.081 | 0.037 | 2.191 | 0.030 |
|  |  |  | Perenniality | 0.001 | 0.043 | 0.035 | 0.973 |
|  |  |  | Woodiness | -0.088 | 0.050 | -1.755 | 0.081 |

Diversification rates tended to be negatively associated with woodiness (Table 2, Fig. 3B), but the relationships were only marginally significant (0.05 < p < 0.10) using the MS and BAMM diversification measures. Relationships between woodiness and Medusa diversification rates, and relationships between perenniality and all diversification rates, were not significant. The standardized partial regression coefficients were similar in magnitude for hybridization metrics and woodiness, while the effects of perenniality were much lower and non-significant (Table 2, Fig. 3B). The best models (as measured by the corrected Akaike Information Criterion, AICc) included either hybridization alone or both hybridization and woodiness, although multiple models fell within two AICc of the best model (Table S7). Multivariate results using alternate parameters, threshold, and priors for estimating diversification rates were qualitatively similar (results not shown).

For completeness, we also examined correlations between hybridization and diversification without accounting for phylogenetic nonindependence. None of these raw correlations were significant, although Medusa diversification rates and HybProp had a trend toward being negative (0.05 < p < 0.10) (Table S8). The estimated relationships with HybProp were all negative, and for HybRatio were all positive, which differed from both the univariate and multivariate PGLS analyses (Fig. 3, Table 1, Table 2), demonstrating the need for phylogenetic context.

### Sister-clade comparisons support a positive hybridization - diversification relationship

We identified 11 pairs of sister families in our dataset with consistent differences in hybridization rates (Table S9). There were no differences in diversification rates estimated by Medusa in any of these pairs (within each pair, Medusa estimated the same net diversification rate for each family), consistent with its recognition of far fewer rate shifts across the phylogeny relative to the other methods. For diversification estimates using both MS and BAMM, there were nine cases where the diversification rates were higher for the family with higher hybridization rates and two cases where the diversification rates were lower for the family with higher hybridization rates. This result was significant using a non-parametric one-tailed sign-test (n = 11, p = 0.033).

## DISCUSSION

### Hybridization and diversification are correlated, but weakly so

The idea that hybridization can promote processes such as adaptation, speciation, and radiation has long been hypothesized, but direct links between hybridization and diversification have been largely untested. Here, we find a positive association between hybridization rates and diversification rates among vascular plant families using three measures of diversification and two measures of hybridization, though these relationships were not always statistically significant, and when significant, models accounted for only a small amount of the observed variation in diversification rates (Fig. 3A, Table 1). Sister-clade comparisons also supported the idea that plant families exhibiting more hybridization tend to diversify faster, as the family with higher hybridization rates also had more rapid diversification (estimated by MS and BAMM) than its sister family in nine of our sample of eleven comparisons (Table S9).

### Potential explanations for relationship between hybridization and diversification

The correlative associations presented here do not necessarily imply that hybridization causes increased diversification. The relationship could be due to causation in either direction or due to correlation with an additional unmeasured or untested variable. Here, we discuss potential mechanisms underlying the association, emphasizing that these mechanisms are speculative and cannot be tested with our observational data.

#### Scenario 1: Increased hybridization leads to faster diversification

Interbreeding between species results in increased genetic and phenotypic diversity which can promote adaptation and speciation. Hybridization can lead to increased genetic diversity (Stebbins 1950, 1959; Hedrick 2013; Suarez-Gonzalez et al. 2018; Marques et al. 2019) and trait diversity, e.g. transgressive segregation (Rieseberg et al. 1999; Seehausen 2004; Kerbs et al. 2017). Hybridization has been shown experimentally to speed up adaptive evolution in annual sunflowers (Mitchell et al. 2019a) and can result in hybrid speciation (for instance in sunflowers, reviewed in Rieseberg 2006) and the Louisiana irises (Arnold et al. 2012), reviewed more broadly in (Mallet 2007; Soltis and Soltis 2009; Arnold et al. 2012; Abbott et al. 2013; Marques et al. 2019). Marques et al. (2019) recently synthesized the idea that the reassembly of old genetic variation into new combinations can provide a substrate for speciation. In contrast to novel mutations arising within a lineage, alleles incorporated from other lineages have previously been tested in other genomic and ecological backgrounds (Mitchell et al. 2019a), and thus may be more likely to promote adaptation. Such adaptation in turn could reduce the chance for extinction and/or promote speciation.

It has also been postulated that hybridization events can lead to adaptive radiation, broadly defined as “the evolution of ecological and phenotypic diversity within a rapidly multiplying lineage”, via similar mechanisms (Seehausen 2004, 2013). For instance, Barrier et al. (1999) infer that the founder of the Hawaiian silversword alliance (an iconic plant radiation) was a hybrid and suggest that enhanced genetic diversity in the founder may have promoted adaptive radiation. Additionally, hybridization can facilitate ecological speciation by providing increased genetic variation and multiple genotypes that can fill available niches, allowing for colonization of novel environments or habitats (Schluter 2000; Seehausen 2004).

In contrast, there is evidence that hybridization could act to decrease diversification. In certain scenarios, hybridization can result in “extinction-by-hybridization” (Campbell et al. In Press; Rhymer and Simberloff 1996; Wolf et al. 2010; Todesco et al. 2016) in which the genetic material of one species is replaced by that of another. At present little data exists to assess how common this process is relative to the above-described processes by which hybridization could enhance diversification.

#### Scenario 2: Faster diversification leads to increased hybridization

Rapid speciation within a lineage results in many closely related species and may also result in incomplete reproductive isolation (e.g., Baldwin and Sanderson 1998; Barrier et al. 1999). Most hybridization takes place within genera (Whitney et al. 2010a), so if fast diversification rates result in many congeneric species then there is enhanced opportunity for hybridization due to the number of potential partners and possibly their geographic proximity. Additionally, taxa that are more closely related have weaker reproductive barriers than distantly related taxa (Coyne and Orr 1989, 1997; Moyle and Nakazato 2010), and these reduced barriers should result in increased interspecific mating (Ellstrand et al. 1996).

#### Scenario 3: Diversification and hybridization are jointly driven by other factors not tested or described here

A correlation between two variables can arise in the absence of a direct causal relationship between them, if they are both driven by a third factor or set of factors. We discuss two of these potential factors (woodiness and perenniality) below.

### Hybridization, diversification, and life history/life form

Previous work has linked both hybridization and diversification to other aspects of plant biology, such as life form and life history. Plant groups with woody growth forms and perennial life histories tend to contain more hybrids (Grant 1958, 1981; Stebbins 1959; Ellstrand et al. 1996; Beddows and Rose 2018; Mitchell et al. 2019b), while plant groups with herbaceous life forms and faster generation times (e.g., annual life histories) have faster diversification (Eriksson and Bremer 1992; Baker et al. 2014) or speciation rates (Dodd et al. 1999). We incorporated perenniality and woodiness into multivariate analyses to ask whether hybridization still had a positive association with diversification rates when these traits are included, and to compare the effect sizes of these potential correlates. We found that hybridization remains positively associated with diversification in these analyses, while woodiness is generally negatively associated with diversification and perenniality had little to no effect (Fig. 3B, Table 2, Table S7). The best model always contained hybridization (Table S7) and the effect of hybridization was always positive and at least weakly significant (Fig. 3B, Table 2). Moreover, the association between hybridization and diversification (measured as the standardized partial regression coefficients) was equivalent to the magnitude of the woodiness – diversification associations, and was stronger than the perenniality – diversification association (Fig. 3B, Table 2). The relationships between woodiness and diversification were in the expected direction (woody groups tended to have lower diversification rates). Interestingly, the best models did not include perenniality (Table S7) and perenniality was not significant (Fig. 3B, Table 2), contrary to expectations (Eriksson and Bremer 1992; Dodd et al. 1999).

### Other correlates of diversification

Although we find detectable associations between hybridization and diversification, our models explain only a small amount of the variation in diversification rates among plants (low adjusted-R^2^ values, Table 1, Table 2). Diversification is an integrative property of a lineage, which is influenced by multiple dimensions of life history, environmental factors, and physiology of the organisms within a lineage. Diversification rates or species richness in plants have been tied to numerous attributes (in addition to life history). These include aspects of the genome such as polyploidy (Vamosi and Dickinson 2006; Wood et al. 2009; Tank et al. 2015; Landis et al. 2018) or genome size (Puttick et al. 2015); dispersal mode or geography such as biotic vs. abiotic dispersal (Larson-Johnson 2016), latitudinal gradients (Davies et al. 2004; Jansson and Davies 2008), biome or habitat differences (Moore and Donoghue 2007; Valente et al. 2010; Goldberg et al. 2011; Onstein et al. 2016), geographic area (Vamosi and Vamosi 2010, 2011), and geographic isolation (Baldwin and Sanderson 1998); species interactions such as defense mutualisms (Weber and Agrawal 2014) and pollination mode (Eriksson and Bremer 1992; Hodges and Arnold 1995; Hodges 1997; Dodd et al. 1999; Vamosi and Vamosi 2010, 2011); and other reproductive aspects such as heterostyly (de Vos et al. 2014), self-incompatibility (Goldberg et al. 2010), and dioecy (Käfer et al. 2014). Future multivariate analyses or phylogenetic path analyses (van der Bijl 2018) with more complete datasets are needed to disentangle and weight the relative contributions of these traits, hopefully resulting in models with greater explanatory power.

### Choice of hybridization and diversification metrics

Our two measures of hybridization (HybProp and HybRatio) and three methods used to estimate diversification rates (MS, Medusa, and BAMM) yield very similar results. Previous work on the correlates of hybridization also found similar results for the two hybridization metrics (Beddows and Rose 2018; Mitchell et al. 2019b). Our results with hybridization propensity and hybrid ratio, though analyses using hybrid ratio tend to show slightly stronger relationships with diversification (Fig. 3, Table 1, Table 2).

There is much debate surrounding the methods of estimation of diversification rates (May and Moore 2016; Moore et al. 2016; Rabosky et al. 2017; Meyer and Wiens 2018; Meyer et al. 2018; Rabosky 2018). For our purposes, the tight correlations among the methods (Fig. 2) and consistency in results across methods (Fig. 3, Table 1, Table 2) suggest a robust relationship between hybridization and diversification. The differences in significance in these methods may be due to the fact that both BAMM and Medusa estimate rate shifts across the phylogeny (Alfaro et al. 2009; Rabosky 2014), meaning that multiple closely-related families can share a diversification rate, while the method-of-moments (MS) estimation is based on branch lengths and richness (Magallon and Sanderson 2001), resulting in a uniquely-estimated diversification rate for each family. Our results suggest that the choice of method to estimate diversification is not crucial to our understanding of the overall direction of relationships between hybridization and diversification, but is important in determining whether the relationship is statistically significant.

### Conclusions

Here, we present evidence that hybridization and diversification rates are positively related across vascular plant families. Although the explanatory power of hybridization is weak, the relationship is detectable even when other factors (perenniality, woodiness) collinear with hybridization rates and previously linked to diversification rates are accounted for. The magnitude of the correlation between diversification and hybridization is on par with that of woodiness and greater than that of perenniality. Although we cannot determine the directionality of causation via data at this taxonomic scale, our results are consistent with experimental evidence indicating that hybridization has the potential to speed up adaptive evolution (Mitchell et al. 2019a), so hybridization may enable more rapid diversification. More likely, hybridization and diversification may have positive effects on each other through a variety of mechanisms. The use of hypothesis-driven phylogenetic path analysis (van der Bijl 2018) may be one way to disentangle causality and incorporate both direct and indirect mechanisms related to diversification rates overall. Detailed evidence suggests that hybridization events may trigger evolutionary radiations, such as in the Hawaiian silverswords (Barrier et al. 1999), Galapagos finches (Grant et al. 2005), Hawaiian crickets (Shaw 2002) or African cichlids (Genner and Turner 2012; Keller et al. 2013; Meier et al. 2017). Exploration of hybridization at the base of additional radiations and incorporation of ancient or cryptic hybridization events are needed to provide further confirmation of this hypothesis. Moreover, a deeper theoretical, empirical, model-based, and experimental knowledge of hybridization, diversification, other factors, and the interplay between them is necessary to more fully understand these evolutionary phenomena.

## Acknowledgments

We thank Jennifer Rudgers, Kellen Paine, Steven Poe, and John Wiens for valuable discussion and feedback on this manuscript, and the UNM Center for Advanced Research Computing for access to supercomputing resources used for the diversification analyses. Funding for the study was provided by NSF DEB 1257965 and UNM (both to K.D.W.).

