## Supplementary Figures and Tables for "Hybridization and diversification are positively correlated across vascular plant families" (from another version of this preprint: version 1, 2019-08-05)

**SUPPLEMENTARY INFORMATION**

**Figure S1.** Distributions and mean values for the number of rate shifts in BAMM using prior expected numbers of 25, 50, 100, and 200.

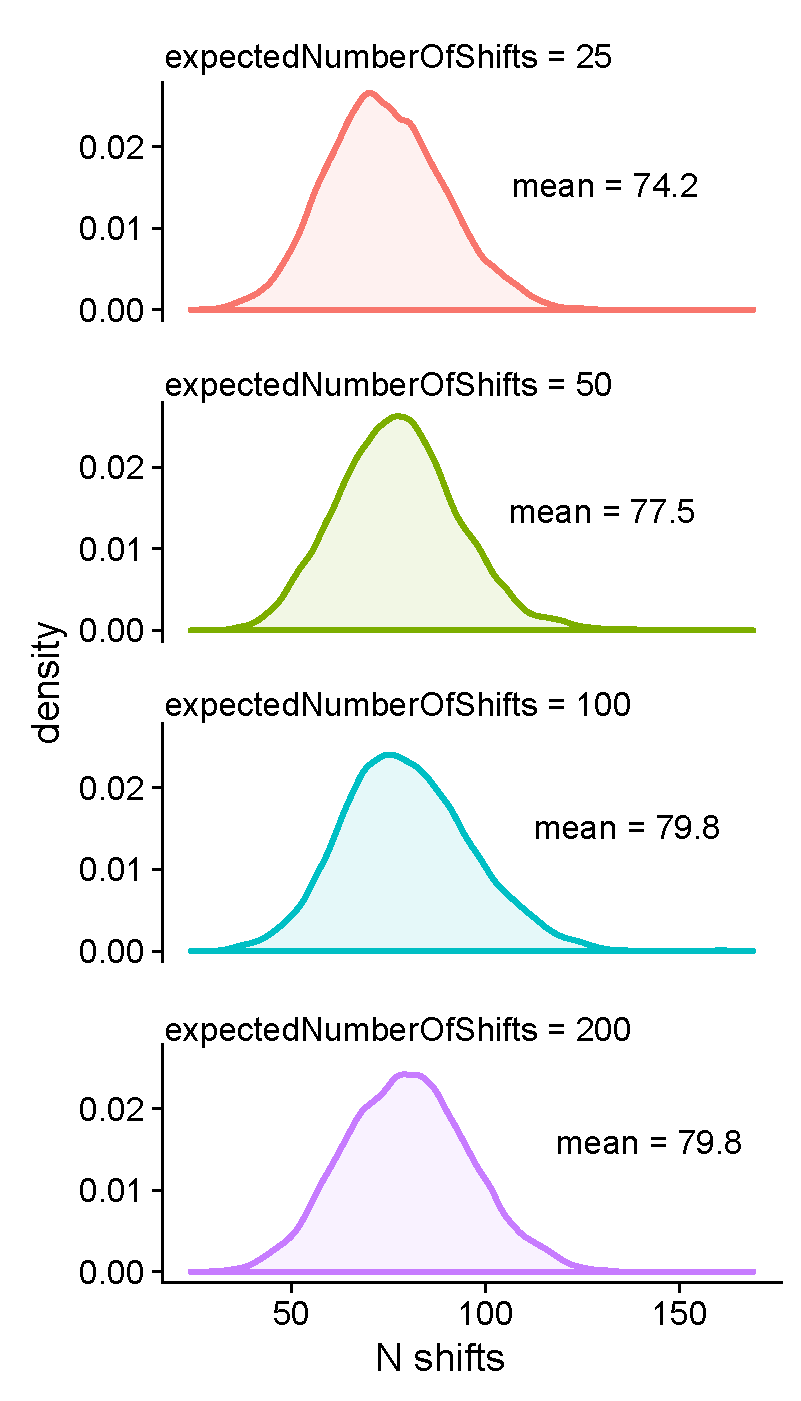

**Table S1.** Method-of-moments (MS) diversification estimates calculated using ϵ = 0.0, 0.5, and 0.9 are significantly correlated. Values below the diagonal are the correlation coefficients, values above the diagonal are the associated p-values.

|  | ϵ = 0.0 | ϵ = 0.5 | ϵ = 0.9 |
| --- | --- | --- | --- |
| ϵ = 0.0 | . | 0.000 | 0.000 |
| ϵ = 0.5 | 0.999 | . | 0.000 |
| ϵ = 0.9 | 0.979 | 0.988 | . |

**Table S2.** Medusa diversification estimates calculated using AICc threshold values of 2, 4, and 8.089 are significantly correlated. Values below the diagonal are the correlation coefficients, values above the diagonal are the associated p-values.

|  | Medusa – 2 | Medusa – 4 | Medusa – 8 |
| --- | --- | --- | --- |
| Medusa – 2 | . | 0.000 | 0.000 |
| Medusa – 4 | 0.995 | . | 0.000 |
| Medusa – 8 | 0.962 | 0.967 | . |

**Table S3.** BAMM diversification estimates estimated using expected number of shifts of 0, 25, 50, and 100 are significantly correlated. Values below the diagonal are the correlation coefficients, values above the diagonal are the associated p-values.

|  | BAMM – 25 | BAMM – 50 | BAMM – 100 | BAMM – 200 |
| --- | --- | --- | --- | --- |
| BAMM – 25 | . | 0.000 | 0.000 | 0.000 |
| BAMM – 50 | 0.999 | . | 0.000 | 0.000 |
| BAMM – 100 | 0.999 | 0.999 | . | 0.000 |
| BAMM – 200 | 0.999 | 0.999 | 1.000 | . |

**Table S4.** Method-of-moments (MS) supplementary results. Univariate results for standardized models of diversification estimates obtained using three different relative extinction fractions (epsilon) for the method-of-moments calculation (ϵ = 0.0, 0.5, and 0.9). Analyses presented in the main text are in bold.

| Epsilon (ϵ) | Hybrid metric | Estimate | Std. Error | T-value | P-value | Adj-R^2^ |
| --- | --- | --- | --- | --- | --- | --- |
| 0.0 | HybProp | 0.110 | 0.065 | 1.683 | 0.094 | 0.009 |
| **0.5** | **HybProp** | **0.116** | **0.066** | **1.754** | **0.081** | **0.011** |
| 0.9 | HybProp | 0.132 | 0.070 | 1.890 | 0.060 | 0.013 |
| 0.0 | HybRatio | 0.124 | 0.056 | 2.237 | 0.026 | 0.020 |
| **0.5** | **HybRatio** | **0.135** | **0.057** | **2.372** | **0.019** | **0.023** |
| 0.9 | HybRatio | 0.161 | 0.061 | 2.644 | 0.009 | 0.030 |

**Table S5**. Medusa supplementary results. Univariate results for diversification estimates obtained using three AICc threshold values: 8, the estimated threshold based on tree size; 5, and 2. These correspond to 49, 70, and 104 shifts, respectively. Analyses presented in the main text are in bold.

| AIC Threshold | Hybrid_metric | Estimate | Std. Error | t value | p-value | Adj-R^2^ |
| --- | --- | --- | --- | --- | --- | --- |
| 8 | HybProp | 0.075 | 0.051 | 1.465 | 0.145 | 0.006 |
| **4** | **HybProp** | **0.102** | **0.055** | **1.859** | **0.065** | **0.013** |
| 2 | HybProp | 0.103 | 0.055 | 1.876 | 0.062 | 0.062 |
| 8 | HybRatio | 0.072 | 0.041 | 1.739 | 0.084 | 0.010 |
| **4** | **HybRatio** | **0.090** | **0.044** | **2.021** | **0.045** | **0.016** |
| 2 | HybRatio | 0.094 | 0.044 | 2.134 | 0.034 | 0.034 |

**Table S6**. BAMM supplementary results. Univariate results for hybridization and BAMM diversification estimates estimated using expected number of shifts of 0, 25, 50, and 100. Analyses presented in the main text are in bold.

| Shift Prior | Hybrid_metric | Estimate | Std. Error | T-value | P-value | Adj-R^2^ |
| --- | --- | --- | --- | --- | --- | --- |
| 25 | HybProp | 0.082 | 0.047 | 1.722 | 0.087 | 0.010 |
| 50 | HybProp | 0.077 | 0.046 | 1.657 | 0.099 | 0.009 |
| **100** | **HybProp** | **0.080** | **0.047** | **1.691** | **0.093** | **0.009** |
| 200 | HybProp | 0.082 | 0.048 | 1.725 | 0.086 | 0.010 |
| 25 | HybRatio | 0.066 | 0.036 | 1.817 | 0.071 | 0.012 |
| 50 | HybRatio | 0.062 | 0.036 | 1.733 | 0.085 | 0.010 |
| **100** | **HybRatio** | **0.066** | **0.036** | **1.817** | **0.071** | **0.012** |
| 200 | HybRatio | 0.067 | 0.036 | 1.835 | 0.068 | 0.012 |

**Table S7.** Comparison of multivariate models. The “best” model for each diversification – hybridization combination(according to AICc) is highlighted in bold.

| Method | Hybrid Metric | Model | AICc | Adj-R^2^ |
| --- | --- | --- | --- | --- |
| MS | HybProp | Hybridization | 470.327 | 0.011 |
| (ϵ = 0.5) |  | Perenniality | 473.376 | -0.005 |
|  |  | Woodiness | 471.340 | 0.005 |
|  |  | Perenniality + Woodiness | 472.552 | 0.004 |
|  |  | Hybridization + Perenniality | 472.360 | 0.006 |
|  |  | **Hybridization + Woodiness** | **469.450** | **0.020** |
|  |  | Hybridization + Perenniality + Woodiness | 470.828 | 0.019 |
|  | HybRatio | Hybridization | 467.783 | 0.023 |
|  |  | Perenniality | 473.376 | -0.005 |
|  |  | Woodiness | 471.340 | 0.005 |
|  |  | Perenniality + Woodiness | 472.552 | 0.004 |
|  |  | Hybridization + Perenniality | 469.747 | 0.019 |
|  |  | **Hybridization + Woodiness** | **466.464** | **0.035** |
|  |  | Hybridization + Perenniality + Woodiness | 467.996 | 0.033 |
| Medusa | HybProp | **Hybridization** | **396.968** | **0.013** |
| (threshold = 4) |  | Perenniality | 400.268 | -0.005 |
|  |  | Woodiness | 400.227 | -0.004 |
|  |  | Perenniality + Woodiness | 402.265 | -0.009 |
|  |  | Hybridization + Perenniality | 398.718 | 0.009 |
|  |  | Hybridization + Woodiness | 398.580 | 0.009 |
|  |  | Hybridization + Perenniality + Woodiness | 400.599 | 0.005 |
|  | HybRatio | **Hybridization** | **396.318** | **0.016** |
|  |  | Perenniality | 400.268 | -0.005 |
|  |  | Woodiness | 400.227 | -0.004 |
|  |  | Perenniality + Woodiness | 402.265 | -0.009 |
|  |  | Hybridization + Perenniality | 397.970 | 0.013 |
|  |  | Hybridization + Woodiness | 397.812 | 0.013 |
|  |  | Hybridization + Perenniality + Woodiness | 399.804 | 0.009 |
| BAMM | HybProp | Hybridization | 346.019 | 0.009 |
| (prior = 100) |  | Perenniality | 348.403 | -0.003 |
|  |  | Woodiness | 346.325 | 0.008 |
|  |  | Perenniality + Woodiness | 348.360 | 0.003 |
|  |  | Hybridization + Perenniality | 347.205 | 0.009 |
|  |  | **Hybridization + Woodiness** | **344.397** | **0.023** |
|  |  | Hybridization + Perenniality + Woodiness | 346.479 | 0.018 |
|  | HybRatio | Hybridization | 345.579 | 0.012 |
|  |  | Perenniality | 348.403 | -0.003 |
|  |  | Woodiness | 346.325 | 0.008 |
|  |  | Perenniality + Woodiness | 348.360 | 0.003 |
|  |  | Hybridization + Perenniality | 346.639 | 0.012 |
|  |  | **Hybridization + Woodiness** | **343.520** | **0.027** |
|  |  | Hybridization + Perenniality + Woodiness | 345.604 | 0.022 |

**Table S8.** Non phylogenetically-informed correlations between diversification rates and hybridization (log-transformed).

| Diversification Method | Hybrid Metric | Correlation | T-value | Degrees of freedom | P-value |
| --- | --- | --- | --- | --- | --- |
| MS | HybProp | -0.097 | -1.36 | 193 | 0.176 |
|  | HybRatio | 0.047 | 0.66 | 193 | 0.510 |
| Medusa | HybProp | -0.120 | -1.68 | 193 | 0.094 |
|  | HybRatio | 0.021 | 0.286 | 193 | 0.775 |
| BAMM | HybProp | -0.070 | -0.97 | 193 | 0.332 |
|  | HybRatio | 0.064 | 0.887 | 193 | 0.376 |

**Table S9.** Sister families used for hybridization and diversification comparisons. Note that all Medusa diversification differences were equal to 0 so they have been omitted.

| Higher Hybridization Family | Lower Hybridization Family | HybProp Difference | HybRatio Difference | MS Difference | BAMM Difference |
| --- | --- | --- | --- | --- | --- |
| Juncaceae | Cyperaceae | 0.044092860 | 0.061506 | -0.050010 | -0.013490 |
| Loasaceae | Hydrangeaceae | 1.111111111 | 0.194444 | -0.003960 | -0.004440 |
| Polygonaceae | Plumbaginaceae | 0.583247014 | 0.098693 | 0.016218 | 0.003414 |
| Brassicaceae | Cleomaceae | 0.265188512 | 0.047160 | 0.013168 | 0.002201 |
| Caprifoliaceae | Adoxaceae | 1.126481734 | 0.045752 | 0.018069 | 0.003867 |
| Tamaricaceae | Frankeniaceae | 0.772834491 | 0.064730 | 0.009212 | 0.001925 |
| Ranunculaceae | Berberidaceae | 0.138248848 | 0.042857 | 0.003923 | 0.000393 |
| Cupressaceae | Taxaceae | 0.738916256 | 0.103448 | 0.001577 | 3.32E-07 |
| Isoetaceae | Selaginellaceae | 0.292917009 | 0.035938 | 0.039802 | 0.020711 |
| Saxifragaceae | Grossulariaceae | 0.908119658 | 0.067308 | 0.010417 | 0.004382 |
| Sarcobataceae | Nyctaginaceae | 1.515151515 | 0.077922 | 0.139177 | 0.030412 |
